# Gene Therapy for Follistatin Mitigates Systemic Metabolic Inflammation and Post-Traumatic Osteoarthritis in High-Fat Diet-Induced Obese Mice

**DOI:** 10.1101/619239

**Authors:** Ruhang Tang, Natalia S. Harasymowicz, Chia-Lung Wu, Kelsey H. Collins, Yun-Rak Choi, Sara J. Oswald, Farshid Guilak

## Abstract

Obesity-associated inflammation and loss of muscle function play critical roles in the development of osteoarthritis (OA); thus, therapies that target muscle tissue may provide novel approaches to restoring metabolic and biomechanical dysfunction associated with obesity. Recent studies indicate that follistatin (FST), a protein which binds myostatin and activin, may have the potential to enhance muscle formation while neutralizing inflammation induced by these proteins. Here, we hypothesized that adeno-associated virus (AAV9) delivery of FST will enhance muscle formation and mitigate metabolic inflammation and knee OA caused by a high fat diet in mice. Obese mice receiving AAV-mediated FST delivery exhibited decreased inflammatory adipokines and cytokines systemically in the serum as well as locally in the joint synovial fluid. Regardless of diet, mice receiving FST gene therapy were protected from post-traumatic OA and bone remodeling induced by joint injury. While obesity disrupted the mitochondrial oxidative phosphorylation (OXPHOS) system in adipocytes, gene therapy for FST restored the key proteins involved in mitochondrial biogenesis, such as PPARγ coactivator 1α and AKT protein kinase 1, leading to the browning of white adipose tissue. Taken together, these findings suggest that FST gene therapy may provide a multifactorial therapeutic approach for injury-induced OA and metabolic inflammation in obesity.

## INTRODUCTION

Osteoarthritis (OA) is a multifactorial family of diseases, characterized by cartilage degeneration, joint inflammation, and bone remodeling. Despite the broad impact of this condition, there are currently no disease-modifying drugs available for OA. Previous studies demonstrate that obesity and dietary fatty acids (FAs) play a critical role in the development of OA, and metabolic dysfunction secondary to obesity is likely to be a primary risk factor for OA (*3*), particularly following joint injury (*1, 2*). Furthermore, both obesity and OA are associated with a rapid loss of muscle integrity and strength (*4*), which may contribute directly and indirectly to the onset and progression of OA (*5*). However, the mechanisms linking obesity, muscle, and OA are not fully understood, and appear to involve interactions among biomechanical, inflammatory, and metabolic factors (*6*). Therefore, strategies that focus on protecting muscle as well as mitigating metabolic inflammation may provide an attractive target for OA therapies in this context.

A few potential interventions, such as weight loss and exercise, have been proposed to reverse the metabolic dysfunction associated with obesity by improving the quantity or quality of skeletal muscle (*7*). Skeletal muscle mass is modulated by myostatin, a member of the transforming growth factor-β (TGF-β) superfamily and a potent negative regulator of muscle growth (*8–10*), and myostatin is up-regulated in obesity and down-regulated by exercise (*11–15*). While exercise and weight loss are the first line of therapy for obesity and OA, several studies involving myostatin delivery have shown small but significant success in achieving long-term maintenance of weight loss or strength gain, particularly in frail or aging populations (*16, 17*). Thus, targeted pharmacologic or genetic inhibition of myostatin provides a promising approach to improving muscle metabolic health by increasing glucose tolerance and enhancing muscle mass in rodents and humans (*10, 18–22*).

Follistatin (FST), a myostatin- and activin-binding protein, has been used as a therapy for several degenerative muscle diseases (*23, 24*), and loss of FST is associated with reduced muscle mass and prenatal death (*25*). In the context of OA, we hypothesize that FST delivery using a gene therapy approach has multifactorial therapeutic potential through its influence on muscle growth via inhibition of myostatin activity (*26*) as well as other members of the TGF-β family. Moreover, FST has been reported to reduce the infiltration of inflammatory cells in the synovial membrane (*27*) and affect bone development (*28*), and pre-treatment with FST has been shown to reduce the severity of carrageenan-induced arthritis (*29*). However, the potential for FST as an OA therapy has not been investigated, especially in exacerbating pathological conditions such as obesity. We hypothesized that overexpression of FST using a gene therapy approach will increase muscle mass and mitigate obesity-associated metabolic inflammation, as well as the progression of OA, in high-fat diet (HFD) induced obese mice.

## RESULTS

### FST gene therapy prevents HFD-induced obesity and inflammation

Dual-energy x-ray absorptiometry (DXA) imaging of mice at 26 weeks of age (Fig. 1A) showed significant effects of FST treatment on body composition. Control-diet, FST-treated mice (i.e., Control-FST mice) exhibited significantly lower body fat percentages, but were significantly heavier than mice treated with a GFP control vector (Control-GFP mice) (Fig. 1B), indicating increased muscle mass rather than fat was developed with FST. With a high-fat diet, control mice (HFD-GFP mice) showed significant increases in weight and body fat percentage that were ameliorated by FST overexpression (HFD-FST mice).

**Figure 1.**
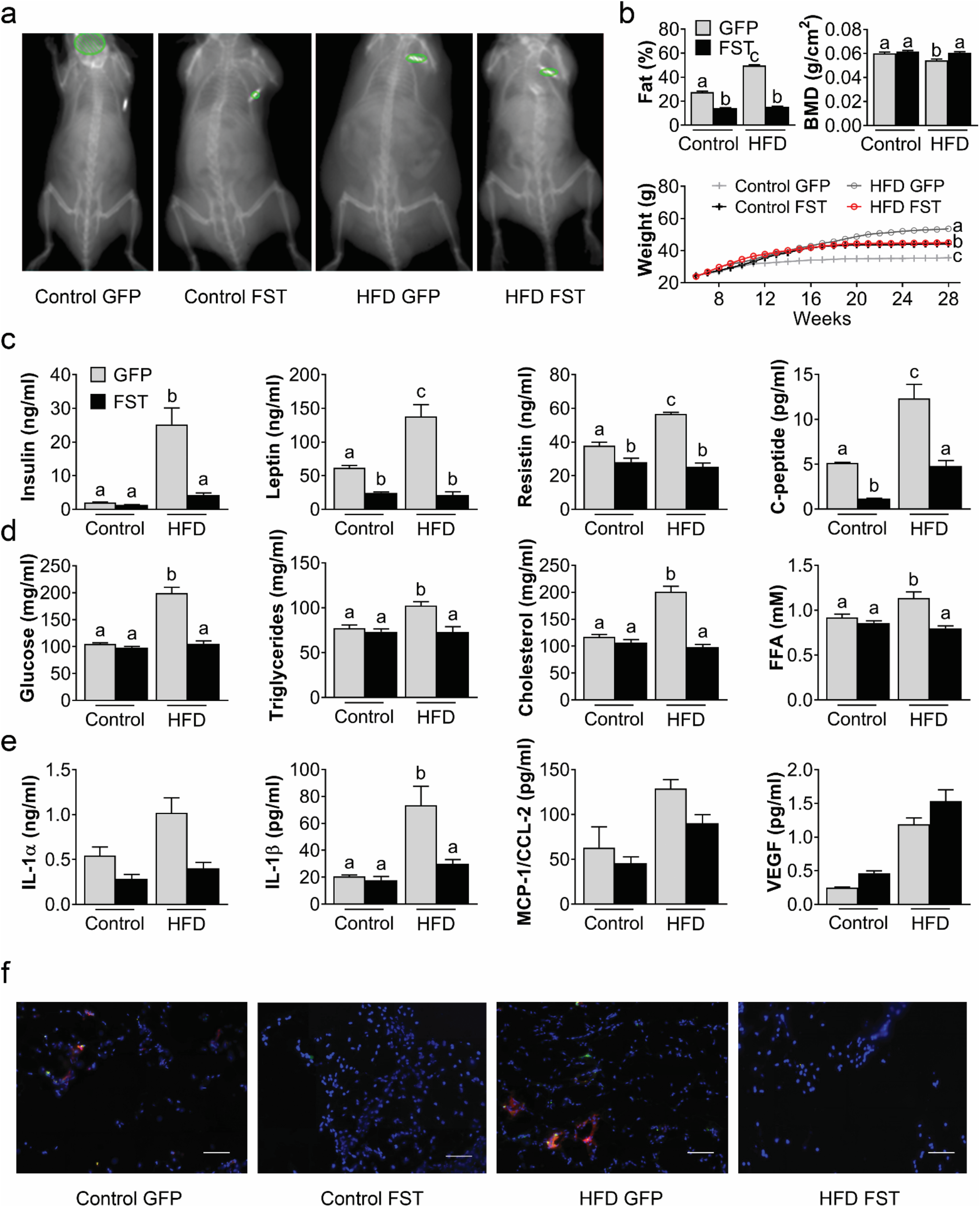
Effects of AAV-mediated FST delivery on body composition and systemic inflammatory markers. (a) Dual-energy x-ray absorptiometry (DXA) images of mice at 26 weeks of age. (b) DXA measurements of body fat percentage and BMD (26 weeks), and body weight measurements over time. (c) Serum levels for adipokines (insulin, leptin, resistin, and C-peptide) at 28 weeks. (d) Metabolite levels for glucose, triglycerides, cholesterol, and free fatty acids (FFAs) at 28 weeks. (e) Serum levels for cytokines (IL-1α, IL-1β, MCP-1, and VEGF) at 28 weeks. (f) Fluorescent microscopy images of visceral adipose tissue with DAPI (blue), CD11b:Alexa Fluor 488 (green), CD11c:PE (red), and DAPI (blue) Scale bar = 100µm. Data presented as mean ± SEM, n=8-10, two-way ANOVA p < 0.05. Groups not sharing the same letter are significantly different with Tukey *post hoc* analysis. For IL-1a and VEGF, p < 0.05 for diet effect and AAV effect. For MCP-1, p < 0.05 for diet effect.

In the high-fat diet group, overexpression of FST significantly decreased serum level of several adipokines including insulin, leptin, resistin, and C-peptide as compared to GFP-treated mice (Fig. 1C). Remarkably, HFD-FST mice also had significantly lower serum levels of glucose, triglycerides, cholesterol, and free fatty acids (FFAs) (Fig. 1D), as well as the inflammatory cytokine IL-1β (Fig. 1E) when compared to HFD-GFP mice. For both dietary groups, AAV-FST delivery significantly increased circulating levels of vascular endothelial growth factor (VEGF), while significantly decreasing IL-1α levels. Furthermore, obesity-induced inflammation in adipose tissue was verified by the presence of CD11b+CD11c+ M1 pro-inflammatory macrophages (Fig. 1F). HFD-FST mice showed decreased occurrence of M1 macrophages in visceral adipose tissue (VAT) compared to HFD-GFP mice.

### FST gene therapy mitigates OA severity and restores muscle performance and pain sensitivity in HFD-fed mice

To determine whether FST gene therapy can mitigate injury-induced OA, mice underwent surgery for destabilization of the medial meniscus (DMM) and were sacrificed 12 weeks post-surgery. Cartilage degeneration was significantly reduced in DMM joints of the mice receiving FST gene therapy in both dietary groups (Fig. 2A, 2C) when compared to GFP controls. FST overexpression also significantly decreased joint synovitis (Fig. 2B, 2D) when compared to GFP controls. To evaluate the local influence of pro-inflammatory cytokines to joint degeneration and inflammation, synovial fluid was harvested from surgical and ipsilateral non-surgical limbs and analyzed using a multiplexed array. The DMM joints from mice with FST overexpression exhibited a trend towards lower levels of pro-inflammatory cytokines, including IL-1α, IL-1β, IL-6, and a higher level of interferon-γ-induced protein (IP-10) in the synovial fluid of DMM joints as compared to contralateral controls (Fig. 2E).

**Figure 2.**
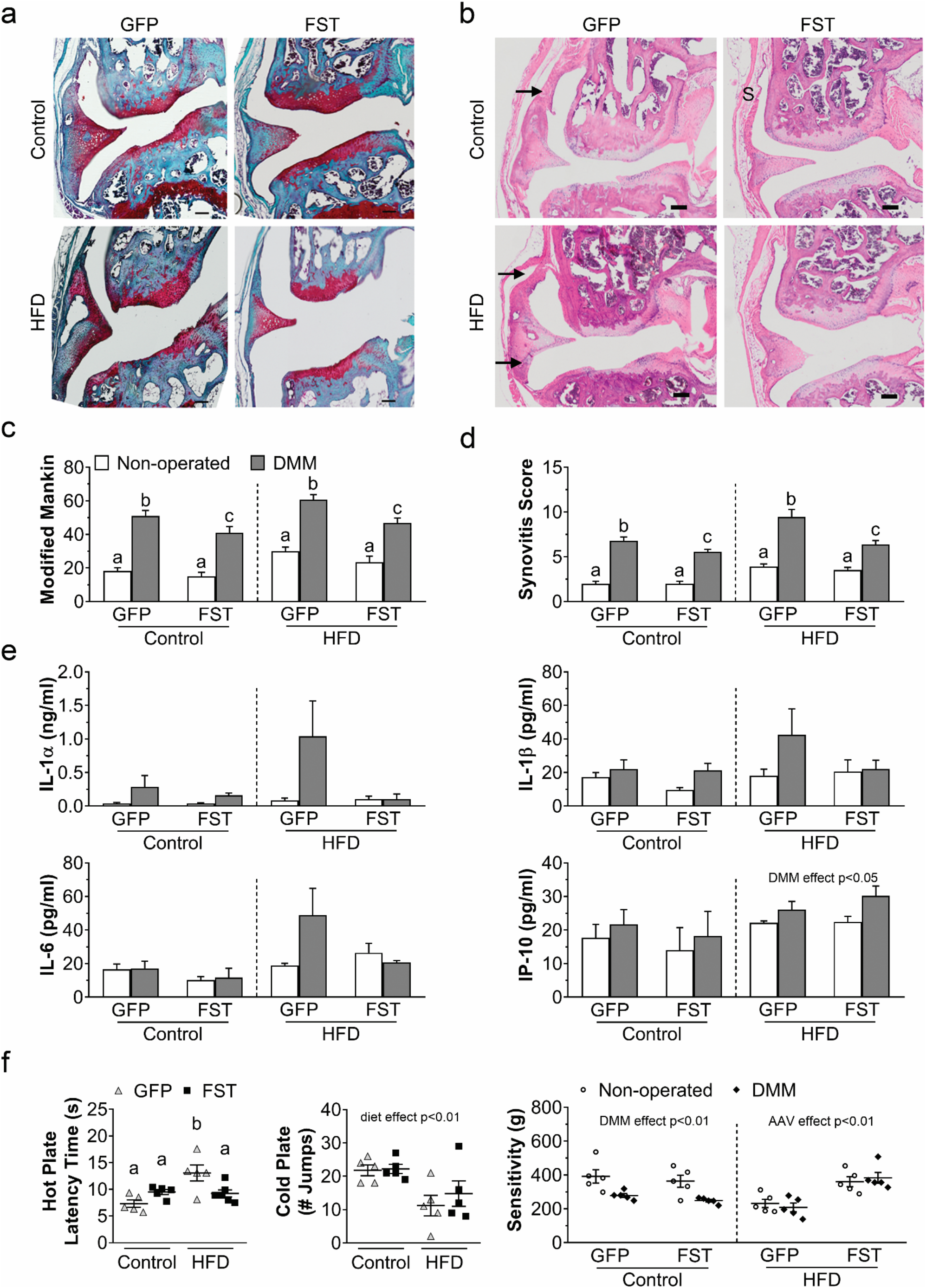
Effects of AAV-FST delivery on OA severity, synovitis, inflammatory cytokines, and pain. (a) Histologic analysis of OA severity via Safranin-O (glycosaminoglycans) and fast green (bone and tendon) staining of DMM-operated joints. (b) Histology (hematoxylin and eosin staining) of the medial femoral condyle of DMM-operated joints (S = synovium). Thickened synovium from HFD mice with a high density of infiltrated cells was observed (arrows). (c) Modified Mankin scores compared within the diet. (d) Synovitis scores compared within the diet. (e) Levels of proinflammatory cytokines in the synovial fluid compared within the diet. (f) Sensitivity to Cold Plate exposure, as measured using the number of jumps in 30 seconds, and Hot Plate latency time, both for non-operated algometry measurements of pain sensitivity compared within the diet. Data presented as mean ± SEM, n=5-10 mice/group, two-way ANOVA p<0.05. Groups not sharing the same letter are significantly different with Tukey *post hoc* analysis.

To investigate the effect of FST on pain sensitivity in OA, animals were subjected to a variety of pain measurements including hot plate, cold plate, and algometry. Obesity increased heat withdrawal latency, which was rescued by FST overexpression (Fig. 2F). Cold sensitivity trended lower with obesity, and since no significant differences in heat withdrawal latency were found with surgery (Supplementary Fig. 1), no cold sensitivity was measured after surgery. Importantly, we found FST treatment protected HFD animals from mechanical algesia at the knee receiving DMM surgery, while Control Diet DMM groups demonstrated increased pain sensitivity following joint injury.

A bilinear regression model was used to elucidate the relationship among OA severity, biomechanical factors and metabolic factors (Supplementary Table 1). Factors significantly correlated with OA were then selected for multivariate regression (Table 1). Both multivariate regression models revealed serum TNF-α levels as a major contributor to OA severity.

**Table 1.** Multivariate regression analyses for factors predicting OA severity. Β = standardized coefficient. ***p < 0.001.

| Predictor Variables | Model 1<br>$\beta$ | Model 2<br>$\beta$ |
| --- | --- | --- |
| Synovial fluid IL-1α | 0.13 | 0.003 |
| Serum IL-1β | 0.006 | 0.073 |
| Serum TNF-α | 0.471 | 0.537 |
| Serum amylin | 0.023 | 0.026 |
| Fat percentage | 0.142 | - |
| Serum leptin | - | 0.00003 |
| Adjusted r<sup>2</sup> | 0.613*** | 0.379*** |

### FST gene therapy enhances muscle growth and restores muscle performance in HFD-fed Mice

We analyzed the effects of FST treatment on muscle structure and mass, and performance measures were conducted on mice in both dietary groups. Both Control-FST and HFD-FST limbs exhibited visibly larger muscles compared to both AAV-GFP groups (Fig. 3A). Additionally, the muscle masses of tibialis anterior, gastrocnemius, and quadriceps each increased significantly with FST treatment (Fig. 3B). Western blot analysis confirmed an increase in FST expression in the muscle at the protein level in FST-treated groups compared to GFP-treated animals in Control and HFD groups (Fig. 3C).

**Figure 3.**
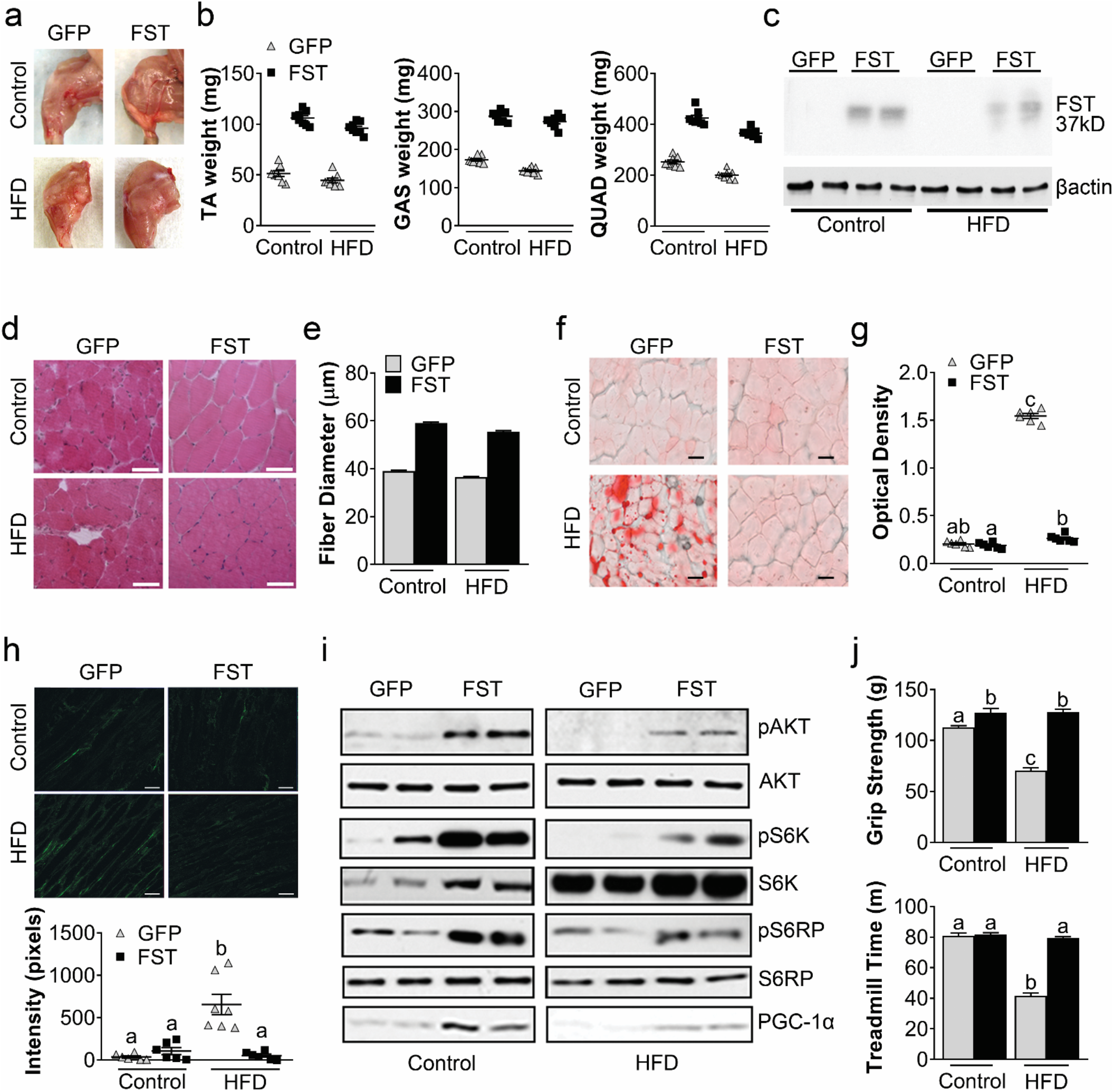
Effects of AAV-FST gene on muscle mass and protein synthesis pathway. (a) Visual and (b) measured mass of tibialis anterior (TA), gastrocnemius (GAS), and quadriceps (QUAD) muscles, n=8, diet, and AAV effects both p<0.05. (c) Western blot showing positive bands of FST protein only in FST-treated muscles, with β-actin as a loading control. (d) H&E stained sections of GAS muscles were measured for (e) mean myofiber diameter, n=100 from 4 mice per group, diet and AAV effects both p<0.05. (f) Oil Red O staining was analyzed for (g) Optical Density values of fatty acids, n=6. (h) Second harmonic generation imaging of collagen in TA sections was quantified for intensity, n=6. (i) Western blotting showing the level of phosphorylation markers of protein synthesis in GAS muscle. (j) Functional analysis of grip strength and treadmill time to exhaustion, n=10. Data presented as mean ± SEM, two-way ANOVA, p < 0.05. Groups not sharing the same letter are significantly different with Tukey *post hoc* analysis.

To determine whether the increases in muscle mass reflected muscle hypertrophy, gastrocnemius muscle fiber diameter was measured in H&E stained sections (Fig. 3D) at 28 weeks of age. Mice with FST overexpression exhibited increased fiber diameter (i.e., increased muscle hypertrophy) relative to the GFP-expressing mice in both diet treatments (Fig. 3E). Oil Red O staining was used to determine the accumulation of neutral lipids in muscle (Fig. 3F). We found that HFD-FST mice were protected from lipid accumulation in muscles compared to HFD-GFP mice (Fig. 3G). Second harmonic generation imaging confirmed the presence of increased collagen content in the muscles of high-fat diet mice, which was prevented by FST gene therapy (Fig. 3H). We also examined the expression and phosphorylation levels of the key proteins responsible for insulin signaling in muscles. We observed increased phosphorylation of Akt^S473^, S6K^T389^ and S6RP-S235/2369 and higher expression of peroxisome proliferator-activated receptor gamma coactivator 1-alpha (*Pgc1a*) in muscles from FST mice compared to GFP mice, regardless of diet (Fig. 3I). In addition to the improvements in muscle structure with HFD, FST overexpressing mice also showed improved function, including higher grip strength and increased treadmill running endurance (Fig. 3J), compared to GFP mice.

Since FST has the potential to influence cardiac muscle as well as skeletal muscle, we performed a detailed evaluation on the effect of FST overexpression on cardiac function. Echocardiography and short-axis images were collected to visualize the left ventricle (LV) movement during diastole and systole (Supplementary Fig. 2A). While the Control-FST mice had comparable LV mass (LVM) and left ventricular posterior wall dimensions (LVPWD) with Control-GFP mice (Supplementary Fig. 3B, 3C), the HFD-FST mice have significantly decreased LVM and trend towards decreased LVPWD compared to HFD-GFP. Interestingly, regardless of the diet treatments, FST overexpression enhanced the rate of heart weight/body weight (HW/BW) (Supplementary Fig. 3D). Although Control-FST mice had slightly increased dimensions of the interventricular septum at diastole (IVSd) compared to Control-GFP (Supplementary Fig. 3E), there was significantly lower IVSd in HFD-FST compared to HFD-GFP. Additionally, we found no difference in Fractial Shortening among all groups (Supplementary Fig. 3F). Finally, transmitral blood flow was investigated using Pulse Doppler. While there was no difference in iso-volumetric relaxation time (IVRT) in Control groups, HFD-FST mice had a moderate decrease in IVRT compared to HFD-GFP. Overall, FST treatment mitigated the changes in diastolic dysfunction and improved the cardiac relaxation caused by HFD.

### FST gene therapy protects trabecular and cortical bone microstructure following injury or HFD

DXA demonstrated that FST gene therapy improved BMD in HFD compared to other groups (Fig. 1B). To determine the effects of injury, diet intervention, and overexpression of FST on bone morphology, knee joints were evaluated by microCT (Fig. 4A). The presence of heterotopic ossification was observed throughout the GFP knee joints, whereas FST groups demonstrated a reduction or an absence of heterotopic ossification. FST overexpression significantly increased the ratio of bone volume to total volume (BV/TV), bone mineral density (BMD), and trabecular number (Tb.N) of the tibial plateau in animals, regardless of diet treatment (Fig. 4B). Joint injury generally decreased bone parameters in the tibial plateau, particularly in control diet mice. In the femoral condyle, BV/TV and Tb.N were significantly increased in mice with FST overexpression in both diet types, while BMD was significantly higher in HFD-FST compared to HFD-GFP mice (Fig. 4B). Furthermore, AAV-FST delivery significantly increased trabecular thickness (Tb.Th) and decreased trabecular space (Tb.Sp) in the femoral condyle of HFD-FST compared to HFD-GFP animals (Supplementary Fig. 3).

**Figure 4.**
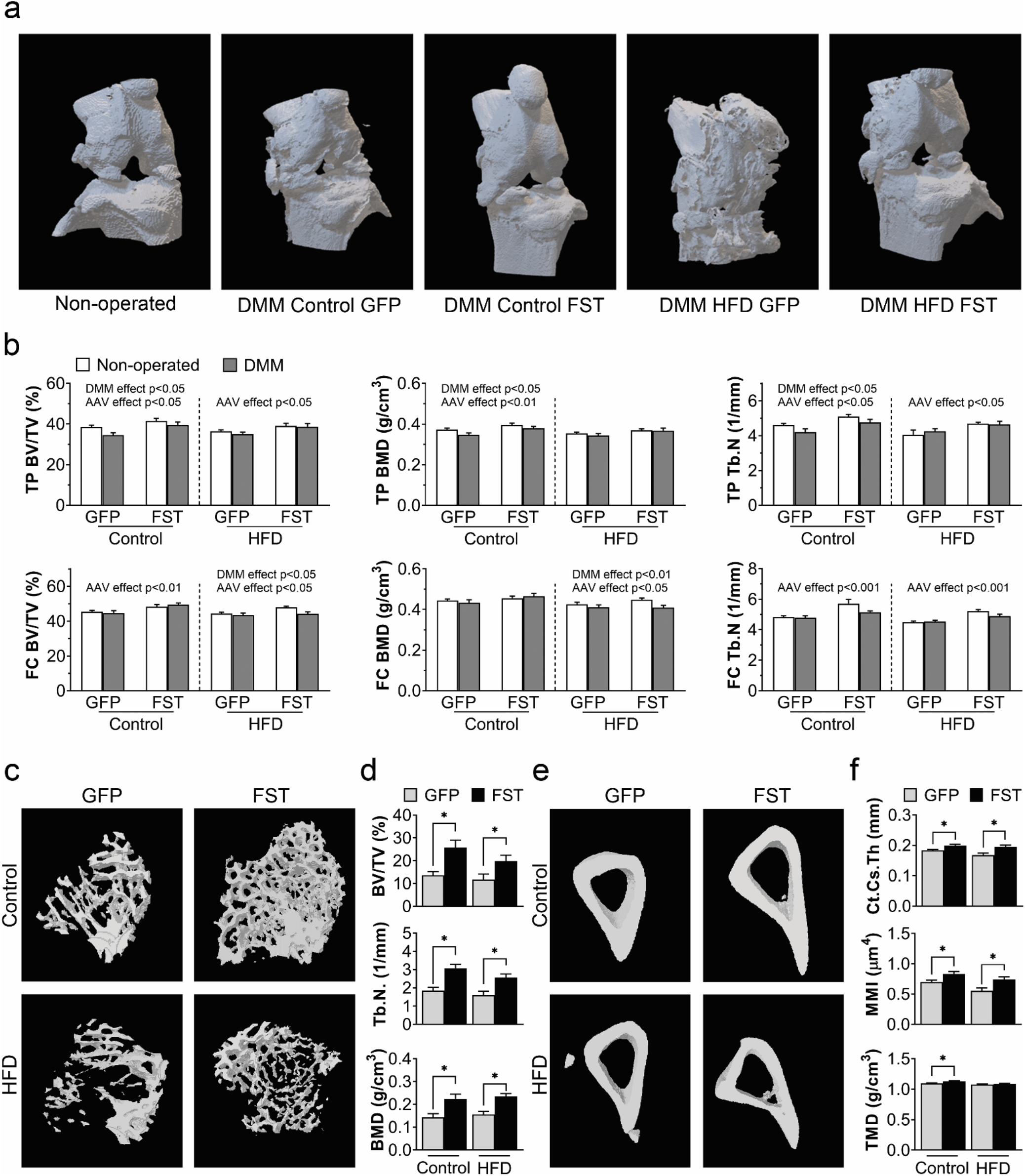
Effects of AAV-FST gene delivery on bone structure. (a) 3D reconstruction of MicroCT images of non-operated and DMM-operated knees. (b) Tibial plateau (TP) and femoral condyle (FC) regional analyses of Trabecular bone fraction bone volume / total volume (BV/TV), bone mineral density (BMD), and trabecular number (Tb.N). Data presented as mean ± SEM, n = 8–19 mice/group, two-way ANOVA. (c) 3D MicroCT reconstruction of metaphysis region of DMM-operated joints. (d) Analysis of metaphysis BV/TV, Tb.N. and BMD. (e) 3D MicroCT reconstruction of cortical region of DMM-operated joints. (f) Analysis of cortical cross-sectional thickness (Ct.Cs.Th), polar moment of inertia (MMI) and tissue mineral density (TMD). (d, f) Data presented as mean ± SEM, n = 8–19 mice/group, Mann-Whitney U test, *p < 0.05. DMM: destabilization of the medial meniscus.

Further microCT analysis was conducted on the trabecular (Fig. 4C) and cortical (Fig. 4E) areas of the metaphyses. FST gene therapy significantly increased BV/TV, Tb.N, and BMD in the metaphyses regardless of the diet (Fig. 4D). Furthermore, FST delivery significantly increased the cortical cross-sectional thickness (Ct.Cs.Th) and polar moment of inertia (MMI) of mice on both diet types, as well as tissue mineral density (TMD) of cortical bones of mice fed Control diet (Fig. 4E).

### FST gene therapy promotes browning of white adipose tissue and rescues mitochondrial oxidative phosphorylation

To elucidate the possible mechanisms by which FST mitigates inflammation, we examined the browning/beiging process in subcutaneous adipose tissue (SAT) with immunohistochemistry (Fig. 5A). Here, we found that key proteins expressed mainly in brown adipose tissue (PGC-1α, PRDM-16, thermogenesis marker UCP-1, and beige adipocyte marker CD137) were up-regulated in SAT of the mice with FST overexpression (Fig. 5B). Increasing evidence suggests that an impaired mitochondrial oxidative phosphorylation (OXPHOS) system in white adipocytes is a hallmark of obesity-associated inflammation (*30*). Therefore, we further examined the mitochondrial respiratory system in SAT. HFD reduced the amount of OXPHOS complex subunits (Fig. 5C). We found that proteins involved in oxidative phosphorylation, including subunits of complex I, II, and III of mitochondria OXPHOS complex, were significantly up-regulated in AAV-FST overexpressing animals compared to AAV-GFP mice (Fig. 5D).

**Figure 5.**
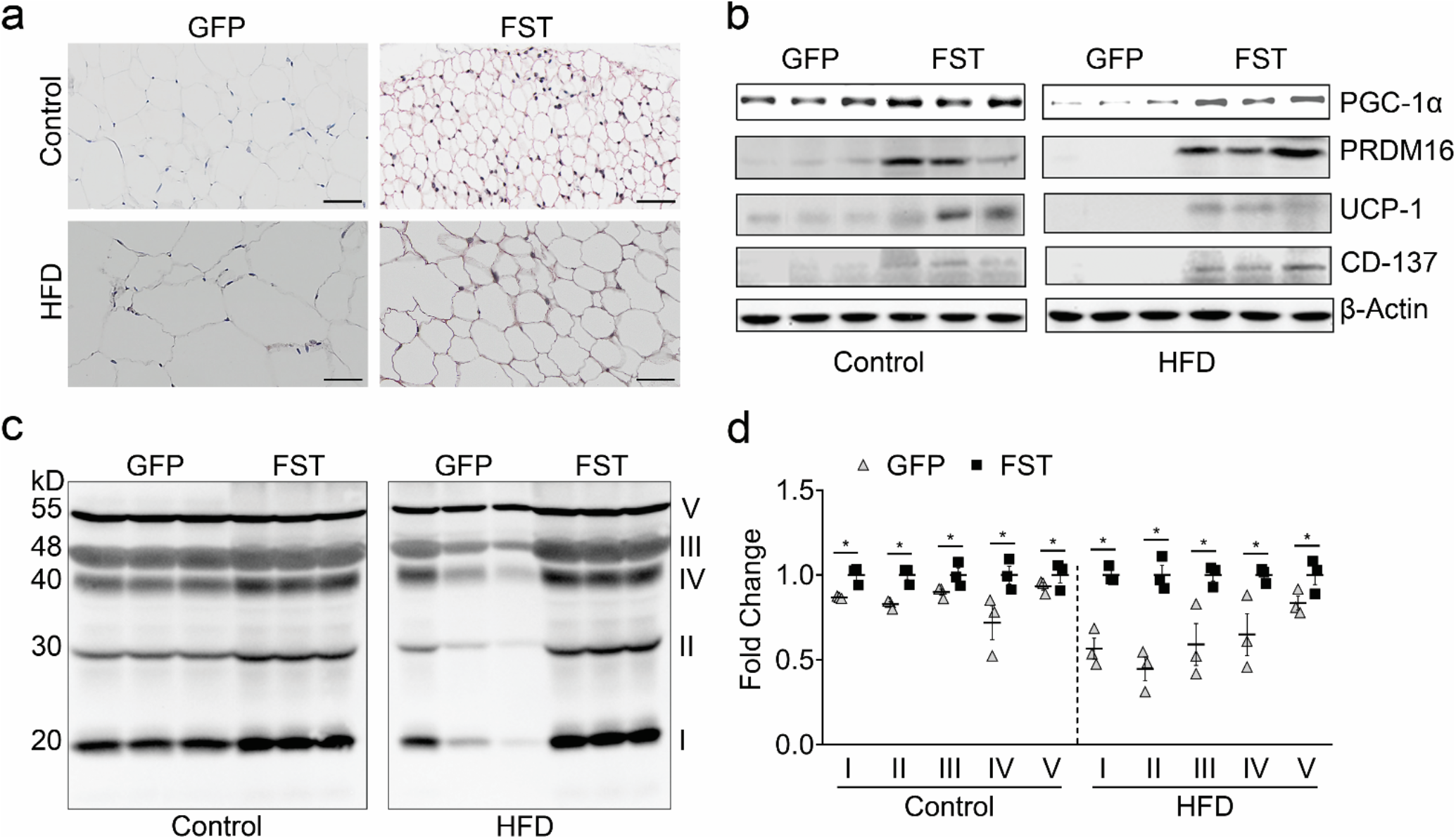
Effects of AAV-FST gene delivery on browning of white adipose tissue and mitochondrial oxidative phosphorylation. (a) Immunohistochemistry of UCP-1 expression in SAT. Scale bar = 50 µm. (b) Western blotting of SAT for key proteins expressed in brown adipose tissue, with β-actin as a loading control. (c) Western blot analysis of mitochondria lysates from SAT for OXPHOS proteins using antibodies against subunits of complex I, II, III and IV, and ATP synthase. (d) Change of densitometry quantification normalized to the average FST level of each OXPHOS subunit. Data presented as mean ± SEM, n=3. *p < 0.05, t-test comparison within each pair.

## DISCUSSION

Our findings demonstrate that a single injection of AAV-mediated FST gene therapy ameliorated systemic metabolic dysfunction and mitigated OA-associated cartilage degeneration, synovial inflammation, and bone remodeling occurring with joint injury and a high-fat diet. Of note, the beneficial effects were observed across multiple tissues of the joint organ system, underscoring the value of this potential treatment strategy. The mechanisms by which obesity and a HFD increase OA severity are complex and multifactorial, involving increased systemic metabolic inflammation, joint instability and loss of muscle strength, and synergistic interactions between local and systemic cytokines (*4, 31*). In this regard, the therapeutic consequences of FST gene therapy also appear to be multifactorial, involving both direct and indirect effects such as increased muscle mass and metabolic activity to counter caloric intake and metabolic dysfunction resulting from a HFD, while also promoting adipose tissue browning. Furthermore, FST may also serve as a direct inhibitor of growth factors in the TGF-β family that may be involved in joint degeneration (*32, 33*).

FST gene therapy showed a myriad of significant beneficial effects on joint degeneration following joint injury while mitigating HFD-induced obesity. These data also indirectly implicate the critical role of muscle integrity in the onset and progression of post-traumatic OA in this model. It is important to note that FST mitigated many of the key negative phenotypic changes previously associated with obesity and OA, including cartilage structural changes as well as bone remodeling, synovitis, muscle fibrosis, and increased pain. Mechanistically, FST restored to control levels a number of OA-associated cytokines and adipokines in the serum as well as the synovial fluid. While the direct effects of FST on chondrocytes remains to be determined, FST has been shown to serve as a regulator of the endochondral ossification process during development (*34*), which may also play a role in OA (*35*). Furthermore, previous studies have shown that a two-week FST treatment of mouse joints is beneficial in reducing infiltration of inflammatory cells into the synovial membrane (*27*). Our findings suggest that FST delivery in skeletally mature mice, preceding obesity-induced OA changes, significantly reduces the probability of tissue damage.

It is well recognized that FST can inhibit the activity of myostatin and activin, both of which are up-regulated in obesity-related modalities and are involved in muscle atrophy, tissue fibrosis and inflammation (*36*). Consistent with previous studies, our results show that FST antagonizes the negative regulation of myostatin in muscle growth, reducing adipose tissue content in animals. Our observation that FST overexpression decreased inflammation at both serum systemic and local joint inflammation may provide mechanistic insights into our findings of mitigated OA severity in HFD-fed mice. Interestingly, our statistical analysis implicated serum TNF-α levels as a major contributor to OA severity, consistent with previous studies linking obesity and OA in mice (*37*). Although the precise molecular mechanisms of FST in modulating inflammation remains unclear, some studies postulate that FST may act like acute phase protein in LPS-induced inflammation (*38*).

In addition to these effects of skeletal muscle, we found that FST gene therapy normalized many of the deleterious changes of a HFD on cardiac function without causing hypertrophy. These findings are consistent with previous studies showing that, during the process of aging, mice exhibiting myostatin KO had an enhanced cardiac stress response (*39*). Furthermore, FST has been shown to regulate activin-induced cardiomyocyte apoptosis (*40*). In the context of this study, it is also important to note that OA has been shown to be a significant risk factor for progression of cardiovascular disease (*41*), and severity of OA disability is associated with significant increases in all-cause mortality and cardiovascular events (*42*).

FST gene therapy also rescued diet- and injury-induced bone remodeling in the femoral condyle, as well as the tibial plateau, metaphysis, and cortical bone of the tibia, suggesting a protective effect of FST on bone homeostasis of mice receiving a HFD. FST is a known inhibitor of BMPs, and thus the interaction between the two proteins plays an essential role during bone development and remodeling. For example, mice grown with FST overexpression via global knock-in exhibited an impaired bone structure (*43*). However, in adult diabetic mice, FST was shown to accelerate bone regeneration by inhibiting myostatin-induced osteoclastogenesis (*44, 45*). Furthermore, it has been reported that FST down-regulates BMP2-driven osteoclast activation (*46*). Therefore, the protective role of FST on obesity-associated bone remodeling, at least in part, may result from the neutralizing capacity of FST on myostatin in obesity. Additionally, improvement in bone quality in FST mice may be explained by their enhanced muscle mass and strength, as muscle mass can dominate the process of skeletal adaptation and conversely, muscle loss correlates with reduced bone quality (*47*).

Our results show that FST delivery mitigated pain sensitivity in OA joints, a critical aspect of clinical OA. Obesity and OA are associated with both chronic pain and pain sensitization (*48*), but it is important to note that structure and pain can be uncoupled in OA (*49*), necessitating the measurement of both behavioral and structural outcomes. Of note, FST treatment protected only HFD animals from mechanical algesia at the knee post-DMM surgery, and also rescued animals from pain sensitization induced by HFD in both the DMM and non-surgical limb. The mitigation in pain sensitivity observed here with FST treatment may also be partially attributed to the antagonistic effect of FST on activin signaling. In addition to its role in promoting tissue fibrosis, activin A has been shown to regulate nociception in a manner dependent on the route of injection (*50–53*). It has been shown that activin can sensitize the TRPV1 channel, leading to acute thermal hyperalgesia (*51*). However, it is also possible that activin and BMP-2 may induce pain indirectly, for example by triggering neuroinflammation (*53–57*), which could lead to sensitization of nociceptors.

The earliest detectable abnormalities in subjects at risk for developing obesity and type 2 diabetes (T2DM) are muscle loss and accumulation of excess lipids in skeletal muscles (*4, 58–61*), accompanied by impairments in nuclear-encoded mitochondrial gene expression and OXPHOS capacity of muscle and adipose tissues (*30, 62–65*). PGC-1α activates mitochondrial biogenesis and increase OXPHOS by increasing the expression of the transcription factors necessary for mitochondrial DNA (mtDNA) replication (*66–68*). We demonstrated that FST delivery can rescue low levels of OXPHOS in HFD diet mice by increasing expression PGC-1α (Fig.3H). It has been reported that high-fat feeding results in decreased PGC-1α and mitochondrial gene expression in skeletal muscles, while exercise increases the expression of PGC-1α in both human and rodent muscles (*61, 69–71*). Although the precise molecular mechanism by which FST promotes PGC-1α expression has not been established, the infusion of lipids decreases expression of PGC-1α and nuclear-encoded mitochondrial genes in muscles (*72*). Thus, decreased lipid accumulation in muscle by FST overexpression may provide a plausible explanation for the restored PGC-1α in the FST mice. These findings were further confirmed by the metabolic profile, showing reduced serum levels of triglycerides, glucose, FFAs, and cholesterol (Fig 1F) and are consistent with previous studies demonstrating muscles with high numbers of mitochondria and oxidative capacity (i.e., Type 1 muscles with high levels of PGC-1α expression) are protected from damage due to a HFD (*4*).

In addition, we found that there is a higher expression level of phosphorylation of protein kinase B (Akt) on Ser473 in skeletal muscle of FST-treated mice in supporting the normal insulin response as compared to untreated HFD counterparts. A number of studies have demonstrated that the serine-threonine protein kinase Akt1 is a critical regulator of cellular hypertrophy, organ size, and insulin signaling (*73*). Muscle hypertrophy both in vitro and in vivo (*74–77*) is stimulated by the expression of constitutively-active Akt1 (*74–77*). In our study, we found that FST overexpression increases the phosphorylation of Akt1. It has been demonstrated that constitutively-active Akt1 also promotes the production of VEGF (*74*) (Supplementary Fig. 2).

Brown adipose tissue (BAT) is thought to be involved in thermogenesis rather than energy storage. BAT is characterized by a number of small multilocular adipocytes containing a large number of mitochondria. The process in which white adipose tissue (WAT) becomes BAT, called “beiging” or “browning”, is postulated to be protective in obesity-related inflammation, as an increase in BAT content positively correlates with increased triglyceride clearance, normalized glucose level, and reduced inflammation. Our study shows that AAV-mediated FST delivery serves as a very promising approach to induce “beiging” of WAT in obesity. A recent study demonstrated that transgenic mice overexpressing FST exhibited an increasing amount of BAT and beigeing in subcutaneous WAT with increased expression of key BAT-related markers including UCP-1 and PRDM16 (*78*). In agreement with previous reports, our data show that *Ucp1*, *Prdm16*, *Pgc1a*, and *Cd167* are significantly up-regulated in SAT of mice overexpressing FST in both dietary interventions. FST has been recently demonstrated to play a crucial role in modulating obesity-induced WAT expansion by inhibiting TGF-β/myostatin signaling and then thus promoting overexpression of these key thermogenesis-related genes. Taken together, these findings suggest that the observed reduction in systemic inflammation in our model may be partially explained by FST-mediated increased process of browning/beiging.

In conclusion, we show that a single injection of AAV-mediated FST, administered after several weeks of HFD feeding, mitigated the severity of OA following joint injury, and improved muscle performance as well as induced beiging of WAT, which together appeared to decrease obesity-associated metabolic inflammation. These findings provide a controlled model for further examining the differential contributions of biomechanical and metabolic factors to the progression of OA with obesity or HFD. As AAV gene therapy shows an excellent safety profile and is currently in clinical trials for a number of conditions, such an approach may allow for the development of therapeutic strategies not only for OA, but also, more broadly, for obesity and associated metabolic conditions.

## MATERIALS AND METHODS

### Animal and Design

All experimental procedures were approved by and conducted in accordance with the Institutional Animal Care and Use Committee guidelines of Washington University in Saint Louis. The overall timeline of the study is shown in Supplementary Fig. 4A. Beginning at 5 weeks of age, C57BL/6J mice (The Jackson Laboratory) were fed either Control or 60% high-fat diet (Research Diets, D12492, n=16/group). At 9 week of age, mice received AAV9-mediated FST or GFP gene delivery via tail vein injection. Destabilization of the medial meniscus (DMM) was used to induce knee OA in the left hind limbs of the mice at the age of 16 weeks. The non-operated right knees were used as contralateral controls. Several behavioral activities were measured during the course of the study. Mice were sacrificed at 28 weeks of age to evaluate OA severity, joint inflammation, and joint bone remodeling.

### Body weight and composition

Mice were weighed bi-weekly. The body fat content and bone mineral density (BMD) of the mice were measured using a dual-energy x-ray absorptiometry scanner (DXA, Lunar PIXImus) at 14 weeks and 26 weeks of age, respectively.

### AAV gene delivery

cDNA synthesis for mouse FST was performed by reverse transcriptase in a RT-qPCR (Invitrogen) mixed with mRNAs isolated from the ovary tissues of C57BL/6J mouse. The PCR product was cloned into the AAV9-vector plasmid (pTR-UF-12.1) under the transcriptional control of the chicken β-actin (CAG) promoter including cytomegalovirus (CMV) enhancers and a large synthetic intron (Supplementary Fig. 4B). Recombinant viral vector stocks were produced at Hope Center Viral Vectors Core (Washington University, St Louis) according to the plasmid co-transfection method and suspension culture. Viral particles were purified and concentrated. The purity of AAV-FST and AAV-GFP were evaluated by SDS-PAGE and stained by Coomassie blue. The results showed that the AAV protein components in 5X 10^11^ vg viral are only stained in three major protein bands: VR1 82kDa, VR2 72kDa, and VR3 62kDa. Vector titers were determined by the DNA dot-blot and PCR methods, and were in the range of 5 × 10^12^ to 1.5 × 10^13^ vector copies (vcs) per ml. AAV was delivered at a final dose of 5 × 10^11^ vg/ mouse by IV tail injection under red light illumination at 9 weeks of age.

### Induction of OA by destabilization of the medial meniscus

At 16 weeks of age, mice underwent surgery for the destabilization of the medial meniscus (DMM) to induce knee OA in the left hindlimb as previously described (*79*). Briefly, anesthetized mice were placed on a custom-designed device, which positioned their hindlimbs in 90-degree flexion. The medial side of the joint capsule was opened and the medial meniscotibial ligament was transected. The joint capsule and subcutaneous layer of the skin were closed with resorbable sutures.

### OA and synovitis assessment

Mice were sacrificed at 28 weeks of age, and changes in joint structure and morphology were assessed using histology. Both hindlimbs were harvested and fixed in 10% neutral buffered formalin (NBF). Limbs were then decalcified in Cal-Ex solution (Fisher Scientific, Pittsburgh, PA, USA), dehydrated and embedded in paraffin. The joint was sectioned in the coronal plane at a thickness of 8 µm. Joint sections were stained with hematoxylin, fast green, and Safranin-O to determine OA severity. Three blinded graders then assessed sections for degenerative changes of joints using a modified Mankin scoring system (*1*). For synovitis, sections were stained with hematoxylin and eosin (H&E) to analyze infiltrated cells and synovial structure. Three independent, blinded graders scored joint sections for synovitis (*1*). Scores for the whole joint were averaged among graders.

### Serum and Synovial Fluid cytokine levels

Serum and synovial fluid (SF) from the DMM and contralateral control limbs were collected as described previously (*1*). For cytokine and adipokine levels in the serum and SF fluid, a multiplexed bead assay (Discovery Luminex 31-plex, Eve Technologies, Calgary, AB, Canada) was used to determine the concentration of Eotaxin, G-CSF, GM-CSF, IFNγ, IL-1α, IL-1β, IL-2, IL-3, IL-4, IL-5, IL-6, IL-7, IL-9, IL-10, IL-12 (p40), IL-12 (p70), IL-13, IL-15, IL-17A, IP-10, KC, LIF, LIX, MCP-1, M-CSF, MIG, MIP-1α, MIP-1β, MIP-2, RANTES, TNF-α, and VEGF. A different kit (Mouse Metabolic Array) was used to measure levels for Amylin, C-Peptide, GIP, GLP-1, ghrelin, glucagon, insulin, leptin, PP, PYY, and resistin. Missing values were imputed using the lowest detectable value for each analyte.

### H&E and Oil Red O staining for muscle

Muscles were cryopreserved by incubation with 2-methylbutane in a steel beaker using liquid nitrogen for 30 seconds, cryoembedded and cryosectioned at 8 µm thickness. Tissue sections were stained following standard H&E protocol. Photomicrographs of skeletal muscle fiber were imaged under brightfield (VS120, Olympus).

Muscle slides fixed in 3.7% formaldehyde were stained with 0.3% Oil red O (in 36% triethyl phosphate) for 30 minutes. Images were taken in brightfield (VS120, Olympus). The relative concentration of lipid was determined by extracting the Oil Red O with isopropanol in equally-sized muscle sections and quantifying the OD_500_ in a 96-well plate.

### Imaging skeletal muscle using second harmonic generation

Second Harmonic Generation (SHG) images of tibialis anterior (TA) were obtained from unstained slices using backscatter signal from an LSM 880 confocal microscope (Zeiss) with Ti:sapphire laser tuned to 820nm (Coherent). The resulting images intensity were analyzed using Image-J software.

### Bone microstructural analysis

To measure bone structural and morphological changes, intact hindlimbs were scanned by microcomputed tomography (microCT, SkyScan 1176, Bruker) with an 18 µm isotropic voxel resolution (455 µA, 700 ms integration time, 4 frame averaging). A 0.5 mm aluminum filter was used to reduce the effects of beam hardening. Images were reconstructed using NRecon software (with 10% beam hardening and 20 ring artifact correction). Subchondral/trabecular and cortical bone regions were segmented using CtAn automatic thresholding software. Tibial epiphysis was selected using the subchondral plate and growth plate as references. Tibial metaphysis was defined as the 1mm region directly below the growth plate. The cortical bone analysis was performed in the mid shaft (4 mm below the growth plate with a height of 1 mm). Hydroxyapatite calibration phantoms were used to calibrate bone density values (mg/cm^3^).

### Immunofluorescent detection of adipose tissue macrophages

Fresh visceral adipose tissues were collected, frozen in OCT, and cryosectioned at 5µm thickness. Tissue slides were then acetone-fixed followed by incubation with Fc-receptor blocking in 2.5% goat serum (Vector Laboratories) and incubation with primary antibodies cocktail containing anti-CD11b:AlexaFluor488 and CD11c:PE (Biolegend). Nuclei were stained with DAPI. Samples were imaged using fluorescence microscopy (VS120, Olympus).

### Histology and Immunohistochemistry for Uncoupling Protein 1

Adipose Tissues were fixed in 10% NBF, paraffin embedded, and cut into 5 µm sections. Sections were deparaffinized, rehydrated and stained with H&E. IHC was performed by incubating sections (n=5 per each group) with the primary antibody (anti-mUCP-1, U6382, Sigma), followed by a secondary antibody conjugated with HRP. Chromogenic substrate 3, 3’-Diaminobenzidine (DAB) was used to develop color. Counterstaining was performed with Harris hematoxylin. Sections were examined under brightfield (VS120, Olympus).

### Western blotting

Proteins of the muscle or fat tissue were extracted using lysis buffer containing 1% Triton X-100, 20 mM Tris-HCL (pH 7.5), 150 mM NaCl, 1 mm EDTA, 5 mM NaF, 2.5 mM sodium pyrophosphate, 1 mM β-glycerophosphate, 1 mM Na_3_VO_4_, 1 µg mL^−1^ leupeptin, 0.1 mM PMSF, and a cocktail of protease inhibitors (Sigma, St Louis, MO, USA, Cat# P0044). Protein concentration were measured with Quick Start^TM^ Bradford Dye Reagent (BIO-RAD). 20 µg of each sample were separated in SDS/PAGE gels with prestained molecular weight markers (BIO-RAD). Proteins were wet-transferred to polyvinylidene fluoride (PVDF) membranes. After incubating for 1.5 h with a buffer containing 5% nonfat milk (BIO-RAD#170-6404) at room temperature in 10 mM Tris-HCl (pH 7.5), 100 mM NaCl and 0.1% Tween-20 (TBST), membranes were further incubated overnight at 4 °C with anti-UCP-1 rabbit polyclonal antibody (1:500, Sigma, U6382), anti-PRDM16 rabbit antibody (Abcam, ab106410), anti-CD137 rabbit polyclonal antibody (1:1000, Abcam, ab203391), total OXPHOS rodent WB antibodies (Abcam, ab110413), anti-Actin (Cell Signaling Technology, 13E5) rabbit mAb (Cell Signaling Technology, 4970), followed by horseradish peroxidase (HRP)-conjugated secondary antibody incubation for 30 min. A chemiluminescent detection substrate (Clarity^TM^, Western ECL) was applied, and the membranes were developed (iBrightCL1000).

### Thermal hyperalgesia experiments

The effects of HFD and FST gene therapy on thermal hyperalgesia were examined at 15 weeks of age. Mice were acclimatized to all equipment one day prior to the onset of testing, as well as a minimum of 30 minutes prior to conducting each test. Thermal pain tests were measured in a room set to 25 °C. Peripheral thermal sensitivity was determined using a hot/cold Analgesia Meter (Harvard Apparatus, Holliston, MA, USA). For hot plate testing, the Analgesia Meter was set to 55 °C. To prevent tissue damage, a maximum cut-off time of 20s was established *a priori* at which time an animal would be removed from the plate in the absence of pain response, defined as paw withdrawal or licking. Animals were tested in the same order 3 times, allowing for each animal to have a minimum of 30 minutes between tests. The analgesia meter was cleaned with 70% ethanol between trials. The average of the three tests was reported per animal. To evaluate tolerance to cold, the analgesia meter was set to 0 °C. After 1h rest, animals were tested for sensitivity to cold over a single 30 second exposure. The number of jumps counted per animal was averaged within each group and compared between groups.

### Mechanical hyperalgesia

Pressure-pain tests were conducted at the knee using a Small Animal Algometer (SMALGO, Bioseb, Pinellas Park, FL, USA). Surgical and nonsurgical animals were evaluated over serial trials on the lateral aspect of the experimental and contralateral knee joints. The average of three trials per limb was calculated for each limb. Within each group, the pain threshold of the DMM limb vs. non-operated limb was compared using a t-test run on absolute values of mechanical pain sensitivity for each limb, p ≤ 0.05.

### Treadmill run to exhaustion

To assess the effect of HFD and AAV-FST treatments on neuromuscular function, treadmill running to exhaustion (EXER3, Columbus Instruments) was performed at 15m/min with 5° inclination angle on the mice 4 month-post gene delivery. Treadmill times were averaged within groups and compared between groups.

### Limb grip test

Forelimb grip strength was measured using Chatillon DFE Digital Force Gauge (Johnson Scale Co.) for front limb strength of the animals. Each mouse was tested 5 times, with a resting period of 90 seconds between each test. Grip strength measurements were averaged within groups and compared between groups.

### Measurement of cardiac function

Cardiac function of the mice at 6 month-old age (n=3) was examined using echocardiography (Vevo 2100 High-Resolution In Vivo Imaging System, VisualSonics) performed by a blinded examiner. Short-axis images were taken to view the left ventricle (LV) movement during diastole and systole. Transmitral blood flow was observed with Pulse Doppler. All data and images were analyzed with an Advanced Cardiovascular Package Software (Visual Sonics).

### Statistical analysis

Detailed Statistical analyses are described in methods of each measurement and its corresponding figure captions. Analyses were performed using GraphPad Prism, with significance reported at the 95% confidence level.

## Acknowledgments

This study was supported in part by NIH grants AR50245, AR48852, AG15768, AR48182, AG46927, AR073752, OD10707, AR060719, AR057235), the Arthritis Foundation, and the Nancy Taylor Foundation for Chronic Diseases.

## Author contributions

RT and FG developed the concept of the study, RT, NSH, CLW, KHC, and YRC collected and analyzed data, SJO analyzed data, and all authors contributed to the writing of the manuscript.

## Data availability statement

All data reported in this paper is available upon request.

---

**Supplementary Table 1.** Correlations of metabolic factors, serum cytokines, synovial fluid cytokines, and joint synovitis with OA severity. *p < 0.05, ***p < 0.001.

| Predictor Variables | r | B (Constant) |
| --- | --- | --- |
| Body weight | 0.306 | 0.586 (23.276) |
| Body gain (week 8 to 28) | 0.269 | 0.486 39.463) |
| SF IL-1α | 0.566*** | 0.008 (45.910) |
| SF IL-1β | 0.266 | 0.128 (45.956) |
| SF IL-6 | 0.598*** | 0.271 (42.745) |
| Serum IL-1α | 0.776*** | 0.023 (36.443) |
| Serum TNF-α | 0.432*** | 1.167 (40.503) |
| Serum VEGF | 0.128 | 2.63 (47.131) |
| Serum resistin | 0.582*** | 0.001 (29.753) |
| Serum C-peptide | 0.566*** | 0.001 (40.804) |
| Serum leptin | 0.562*** | 0* (41.526) |
| Serum amylin | 0.464*** | 0.064 (40.649) |
| Serum insulin | 0.398* | 0 (45.997) |
| Percent fat | 0.641*** | 0.553 (34.569) |
| Synovitis | 0.749*** | 4.28 (19.746) |

**Supplementary Figure 1.**
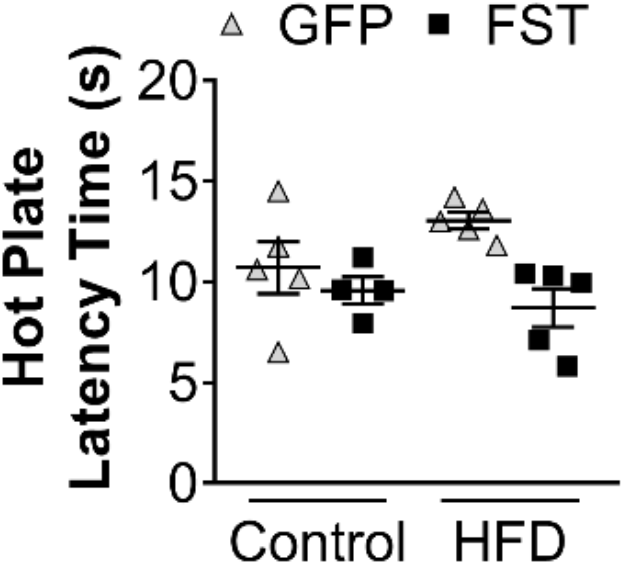
Hot Plate latency time for DMM-operated. Data presented as mean ± SEM, n=4-5 mice/group. No significant differences were observed among the different groups.

**Supplementary Figure 2.**
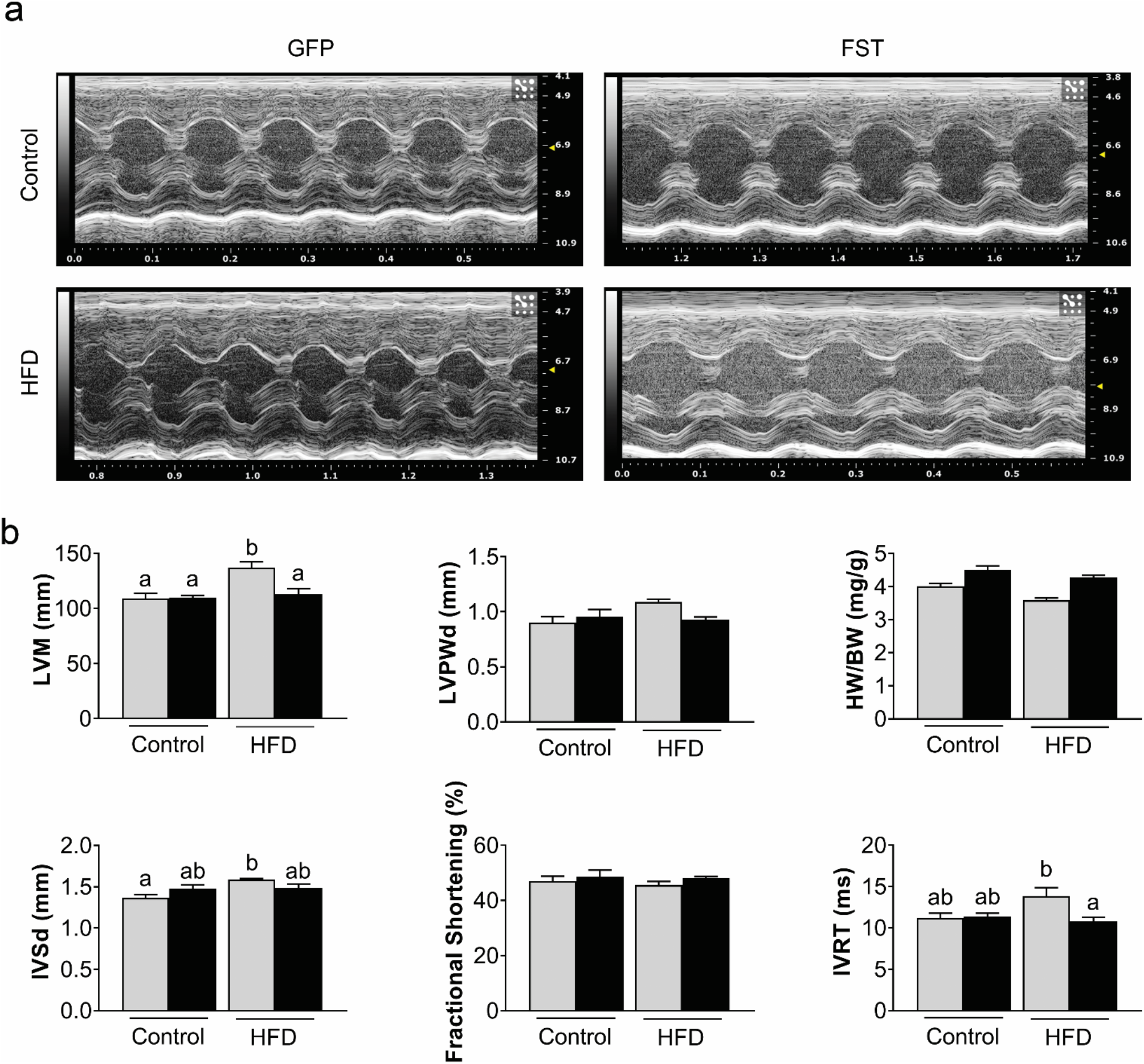
Cardiac structure and function analyzed by echocardiography. (a) Representative two-dimensional short axis views and M-mode images, while images were taken to view the left ventricle (LV) movement during diastole and systole. (b) LV mass (LVM), left ventricular posterior wall dimensions (LVPWd), heart weight/body weight (HW/BW), dimensions of the interventricular septum at diastole (IVSd), % Fractional Shortening, and left ventricular iso-volumetric relaxation time (IVRT). Data presented as mean ± SEM, two-way ANOVA, p < 0.05, n = 3. Groups not sharing the same letter are significantly different with Tukey *post hoc* analysis. Plots not indicating stats showed statistical significance p<0.05 only for HW/BW diet effect and AAV effect.

**Supplementary Figure 3.**
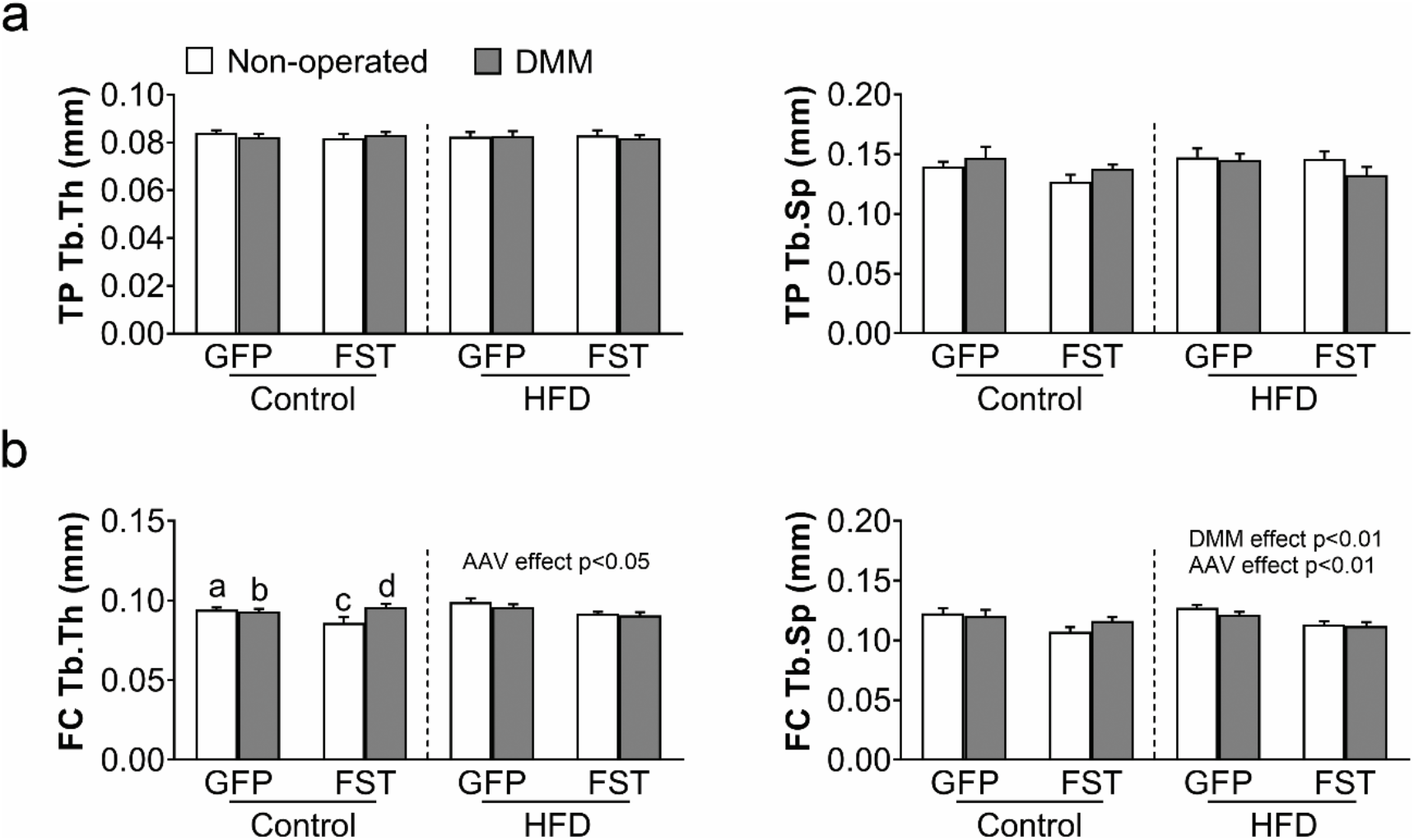
Additional microCT data. (a) Tibial plateau (TP) region analysis of Trabecular thickness (Tb.Th) and Trabecular spacing (Tb.Sp). (b) Femoral condyle (FC) region analysis of Tb.Th and Tb.Sp. Data presented as mean ± SEM, n = 8–12 mice/group, two-way ANOVA p < 0.05. Groups not sharing the same letter are significantly different with Tukey *post hoc* analysis.

**Supplementary Figure 4.**
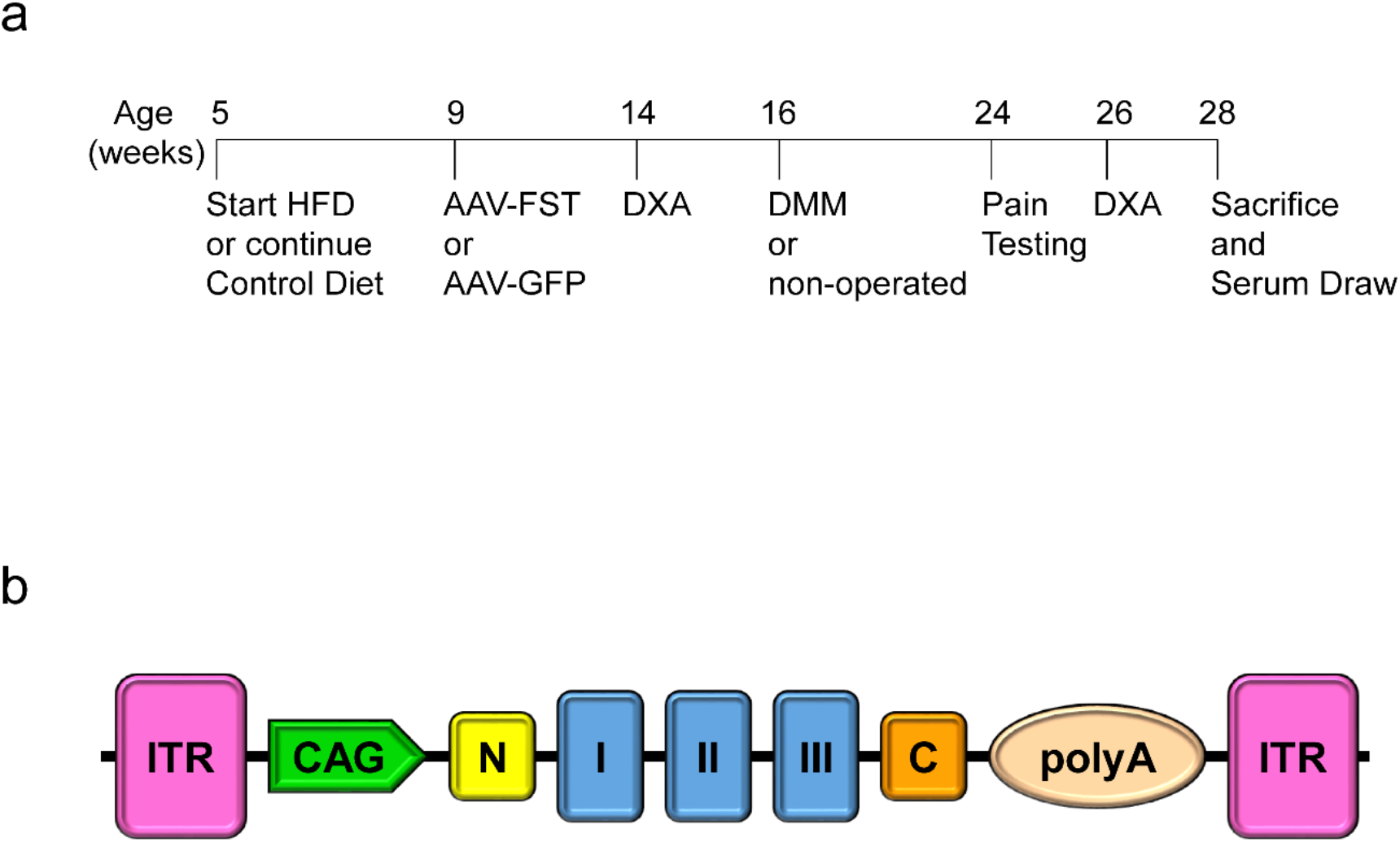
(a) Timeline of major experimental outcomes. (b) AAV vector.

